# Adaptive *P*-element insertions in a long non-coding RNA are an emerging piRNA cluster

**DOI:** 10.1101/2025.07.17.665384

**Authors:** Savana Hadjipanteli, Natalie Copeland, Nancy Marmolejo Bustamante, Maureen Nweke, Basabdatta Adhikari, Erin S. Kelleher

## Abstract

Transposable elements are genetic parasites whose mobilization throughout the genome is a major source of deleterious mutations. However, some TE insertions are beneficial because they improve host fitness. Adaptive TE insertions sometimes alter the function of adjacent genes by positively and negatively impacting their expression, or by altering their encoding proteins. Alternatively, individual TE insertions can also be adaptive because they occur in piRNA clusters and lead to piRNA-mediated silencing of transposition. In a recent laboratory evolution experiment, we discovered that the long non-coding RNA *CR43651* is an adaptive insertion hot spot for *P*-element DNA transposons in *Drosophila melanogaster*. The functional effects of these insertions on *P*-element repression and *CR43651* function were unknown. In this study, we examined the effects of *CR43651* insertions on *P*-element transcriptional regulation, piRNA biogenesis, and viability. We determined that although *CR43651* is not a canonical piRNA cluster, chromosomes containing antisense *P-*element insertions in *CR43651* exhibit enhanced piRNA-like silencing in the stage of oogenesis when *P-*elements transpose, potentially explaining their adaptive benefit. We also discovered that the fitness benefit provided by *P-*element repression is offset by recessive viability effects of insertion chromosomes, potentially due to disrupted production of *mir14*, a miRNA produced from *CR43651*.

## INTRODUCTION

Transposable elements (TEs) are mobile genetic parasites that replicate in germline cells. Through transposition, they cause DNA damage and produce new mutations, both of which are harmful to host fitness. In eukaryotes, piRNA clusters play a key role in defending host genomes from these fitness costs by producing the Piwi-interacting RNAs (piRNAs, 23-32 nt) that regulate TE activity and genomic spread. These specialized loci often contain insertions from multiple TE families, which are transcribed and processed into diverse piRNA species (Mével-Ninio et al. 2007; Brennecke et al. 2007; reviewed in Le Thomas, Tóth, et al. 2014). piRNA then enact co-transcriptional and post-transcriptional regulation of genomic targets, including homologous TE loci and TE-derived mRNA (Senti et al. 2015; Yu et al. 2015).

TEs often invade the genome of a new host through horizontal transfer between non-mating species. These newly invading TEs pose a challenge to host defenses because they are not targeted by the existing pool of piRNA. For invading TEs, piRNA clusters are proposed to act as a genetic “trap” such that random transposition of an invading TE copy into a piRNA cluster leads to piRNA production and silencing of all TE copies in *trans* (Bergman et al. 2006; Brennecke et al. 2007; Zanni et al. 2013; Kelleher et al. 2018; Kofler 2019). However, transposition into the trap may only be the first stage of piRNA mediating silencing of a TE. Once silencing is initiated, TE insertions outside of major piRNA clusters are converted into piRNA producing sites (Shpiz et al. 2014; Mohn et al. 2014). Furthermore, for many TE families, stand-alone insertions outside of large piRNA clusters are proposed to act as the major source of piRNA, with large piRNA clusters being dispensable for TE control (Gebert et al. 2021).

Hence it is proposed that once TEs fall into the trap, new piRNA clusters emerge and over time, major piRNA clusters may become lost (Srivastav et al. 2024; Gebert et al. 2021) *P-*elements are DNA transposons that telomeric associated sequences (TAS) natural populations of *Drosophila melanogaster*. Observations of *P-*element repression have been foundational in revealing piRNA mediating silencing and motivating the “trap” model. Genetic mapping of *P-*element repression revealed that insertions in particular loci have the capacity to regulate *P-*element transposition (Ronsseray et al. 1996; Marin et al. 2000; Stuart et al. 2002)., These loci were later revealed to be piRNA clusters located in the telomere-associated sequences (TAS) (Karpen & Spradling 1992; Brennecke et al. 2007; reviewed in Kelleher 2016). In North America—the epicenter of the *P-*element invasion—the overwhelming majority of genomes contain *P-*element insertions in TAS (Zhang et al. 2020; Stuart et al. 2002).

We recently recreated the *P-*element invasion into a single naive genome (DSPR founder A4; King et al. 2012), and maintained 10 replicate invasion populations over a three year experiment. Consistent with observations from natural populations: 9 out of 10 laboratory populations rapidly evolved piRNA mediated repression that was coincident with insertions into TAS piRNA clusters (Wang et al. 2023). However, we discovered that these same populations contained numerous independent *P-*element insertions into a specific long non-coding RNA locus: *CR43651*. These insertion alleles were often segregating at elevated frequencies in experimental populations, indicating they confer a selective advantage. Furthermore, they lead to modest piRNA production from flanking genomic DNA in *CR43651*, suggesting they cause *CR43651* to act as a piRNA source locus in at least a subset of ovarian cells. However the contribution of *CR43651* insertion alleles to *P-*element control remained unclear.

In this study, we examined the silencing properties and viability effects of chromosomes containing *P*-element insertions into *CR43651*. We discovered that surprisingly, two insertion chromosomes containing an antisense *P-*element insertion in *CR43651* exhibit enhanced piRNA production and silencing of homologous reporters in *trans. Trans-*silencing in particular occurs even in the absence of *P-*elements in TAS, and is more robust than TAS-dependent silencing at the earliest stages of oogenesis when *P-*elements transpose. Unlike TAS piRNA clusters however, *CR43651* is not a “trap”, as insertions of exogenous sequences into *CR43651* alone are not sufficient to establish piRNA silencing. We propose that *CR43651* insertions may represent an emerging piRNA cluster, which over time could acquire fully autonomous properties.

## RESULTS

### *P-*element insertions in *CR43651* are piRNA source loci

To isolate chromosomes containing *P*-element insertions in *CR43651*, we generated second chromosome extraction lines from two experimental populations from Wang et al (2023), (populations 10 and 25), in a repressive *TP5* background (Stuart et al. 2002). *TP5* is a *P-* element insertion in the X chromosome TAS piRNA cluster (*X*-TAS), which establishes piRNA mediated silencing of *P-*elements *in trans*. By generating chromosome extractions in a *TP5* background, we ensured that *P-*elements in balanced extraction lines (*TP5/FM6;insertion chromosome/CyO; +)* were transpositionally stable and new insertions were not produced. We identified extraction chromosomes containing *CR43651* insertions via PCR, and obtained three such chromosomes in a balanced *TP5* background: *S10-3, S10-7* and *S25-3. S10-3* and *S10-7* contain the same antisense insertion (A4 2R:9279721-) whereas *S25-3* contains a different sense insertion (A4 2R:9279848+).

We further performed deep sequencing of *S10-3, S10-7* and *S25-3* genotypes to uncover additional *P-*element insertions carried on these chromosomes. The *S10-3, S10-7* and *S25-3* genotypes contain 6,5, and 7 additional *P-*element insertions, along with their known insertions in *CR43651* and *X-*TAS (Supplementary Table 1, Figure 1A). Consistent with their origins as second chromosome extraction lines, all insertions occurred on the second chromosome with the exception of *TP5* and a single rare insertion on chromosome 3L in *TP5/FM6;S10-3/CyO* (Figure 1A, Supplementary Table 1). We also discovered that *S10-3* and *S10-7* share two additional *P-*element insertions on 2L, in addition to the antisense *P-*element in *CR43651*. Notably, outside of *TP5*, no *P-*element insertions occur in annotated ovarian piRNA clusters in the A4 genetic background, although one insertion from *S10-3* occurs in a region that was identified as a source of piRNAs in A4 ovaries only one of two biological replicates (Supplementary Table 1; Srivastav et al. 2024).

**Figure 1.**
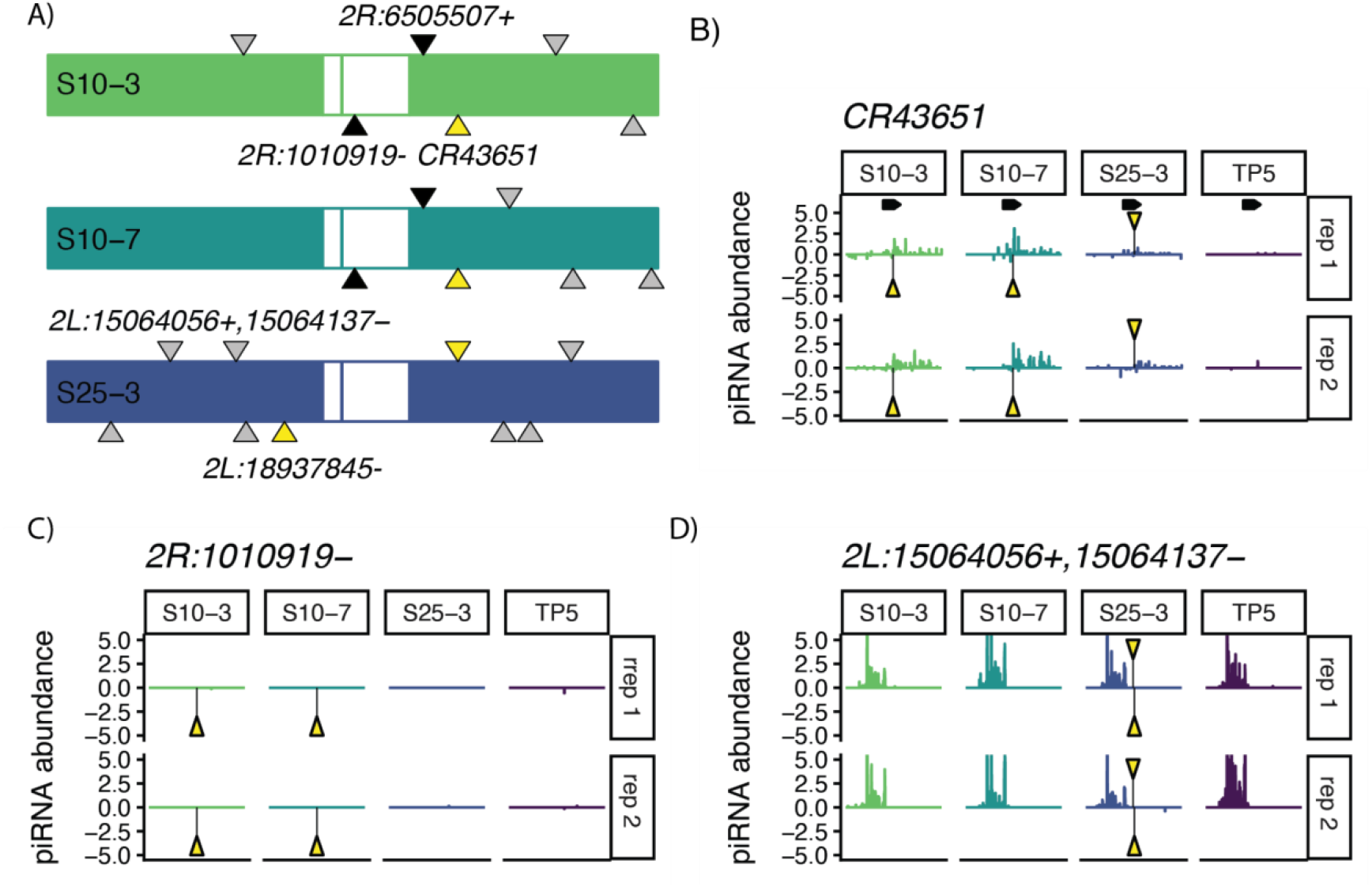
A) Second chromosome maps of *S10-3, S10-7* and *S25-3*. Location of *P* element insertions are indicated by triangles. Yellow triangles highlight insertions that act as *de novo* piRNA clusters, black triangles indicate two additional insertions shared between *S10-3* and *S10-7*. A single low frequency insertion on 3L in *S10-3* is not shown. B-D) Sliding window analysis of uniquely mapping piRNAs aligning to the 6 Kb window surrounding insertion site shows 3 representative patterns. B) Insertions in *CR43651* show piRNA production from both genomic strands surrounding the insertion site. C and D) Insertions in other locations have no obvious impact on piRNA production, as piRNAs are absent (C) or present and unaltered (D) regardless of the insertion status. Genome and small RNA sequencing underlying all panels are in PRJNA1292797.

Euchromatic TEs that are targets of piRNA silencing are often converted to piRNA source loci, leading to piRNA production from DNA adjacent to the insertion site (Shpiz et al. 2014; Mohn et al. 2014). The specific requirements for this conversion are not fully understood; however, it appears that an intact TE promoter is required (Olovnikov et al. 2013; Shpiz et al. 2014). Since the insertions on extraction chromosomes were very recently produced, and the promoter of *P*-elements are not frequently mutated during transposition (Engels et al. 1990; Kaufman & Rio 1991), the majority of the insertions in the genotypes we study here likely contain intact promoters. To examine whether these insertions act as piRNA source loci, we examined the piRNA pools of *TP5* and *TP5;insertion chromosome* homozygotes (Figure 1B-D). *CR43651* is not a piRNA cluster, and no piRNA biogenesis is observed from this locus in the *TP5* background. However, consistent with our previous results from pooled populations samples (Wang et al. 2023), *P-*element insertion alleles in *CR43651* exhibit piRNA biogenesis from opposing genomic strands that spread from the insertion site (Figure 1B). Our chromosome extraction lines further reveal that significantly more piRNA production is observed surrounding the antisense insertion in *S10-3* and *S10-7*, where transcription from the *P-*element promoter would be convergent with *CR43651*.

Interestingly, we observed that most other *P-*element insertions on extraction chromosomes did not exhibit the same behavior as *CR43651* in promoting piRNA biogenesis from surrounding genomic DNA. Of 16 other insertions we uncovered at various genomic locations, only one additional insertion was associated with *de novo* piRNA production from adjacent genomic sites (*2L:18937845-* on the S25-3 chromosome, Figure 1A, Supplementary Figure 1). The 6Kb regions surrounding all other insertion sites were either quiescent with regards to piRNA production, or exhibited piRNA production that was unaltered by *P-*element insertions (Figure 1C,D Supplementary Figure 1). Thus while an intact promoter might be required to convert piRNA targets to piRNA source loci, our observations suggest additional unknown requirements in *de novo* piRNA cluster formation.

### *CR43651* insertions chromosomes enhance *P-*element derived piRNA production

We next considered how insertion chromosomes impact *P-*element derived piRNA production. Additional *P-*element copies are predicted to amplify *P-*element derived piRNA production, both through sense *P-*element mRNAs that feed forward piRNA production through the ping-pong amplification loop (Brennecke et al. 2008; Kelleher & Barbash 2013), and through the conversion of additional *P-element* insertions into *de novo* piRNA clusters (Olovnikov et al. 2013). Consistent with this, we observed that *P-*element derived piRNAs were dramatically increased in genotypes containing insertion chromosomes (Figure 2A,B). We did not observe a systematic increase in piRNAs from other TE families in association with insertion chromosomes (Figure 2C). *P-*element piRNA production is amplified more in the *S10-3* and *S10-7* containing genotypes, where antisense insertions into *CR43651* establish robust piRNA production (Figure 1B). Notably, the difference in *P-*element piRNA production between the insertion chromosomes is not related to copy number: as *S25-3* contains the most *P-*element copies but exhibits the lowest increase in *P-*element derived piRNA abundance (Figure 1A, Figure 2B,C).

**Figure 2.**
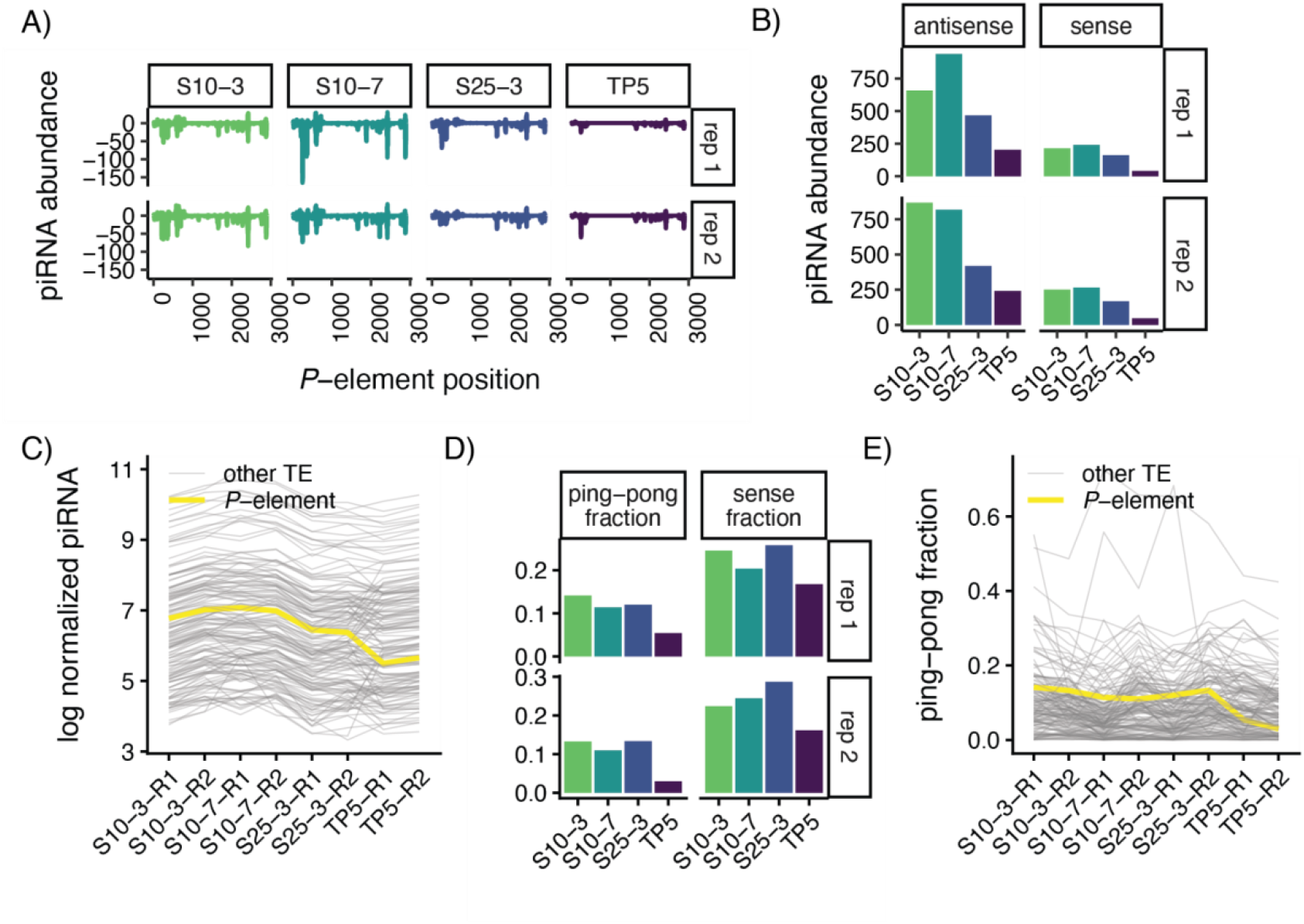
Insertion chromosomes amplify *P*-element derived piRNA production. A) Sliding window analysis of piRNAs (23-32 nt) mapping to *P*-elements. B) Abundance of piRNAs from sense and antisense *P*-element strand C) Normalized abundance of *P*-element derived piRNA in all samples as compared to all other TE families. D)ping-pong fraction and sense piRNA fraction of *P*-element derived piRNAs. E) Ping-pong fraction of *P*-element derived piRNA in all samples as compared to all other TE families, small RNA sequencing underlying all panels are in PRJNA1292797.

We further investigated how ping-pong amplification of *P-*element derived piRNA was altered by insertion chromosomes. Consistent with *P-*element mRNA acting to feed forward piRNA production through ping-pong amplification, we observed that the ping-pong fraction and the sense fraction of *P*-element piRNAs was increased in all three insertion chromosomes (Figure 2D,E). This change was also unique to *P-*elements as no other TE family exhibited a systematic change in the ping-pong signature in the presence of insertion chromosomes (Figure 2E). However, increased ping-pong alone does not appear to explain the disproportionate increase of *P-*element piRNA in association with *S10-3* and *S10-7*. While the ping-pong fractions and antisense fractions are similar for all three insertion chromosomes, the overall piRNA abundance increases a lot more in the presence of *S10-3* and *S10-7* (Figure 2A-2E). This suggests that *S10-3* and *S10-7* must have additional effects on piRNA biogenesis beyond feeding forward ping-pong amplification.

### *trans* silencing in association with *CR43651* extraction chromosomes

We next evaluated the silencing capacity of insertion chromosomes in the female germline using *P{lacZ}* reporters (O’Kane & Gehring 1987). These reporter genes are regulated by the *P-*element promoter and share 503 bp of expressed homologous sequence with native *P-* elements, therefore, their expression is sensitive to the production of *P-*element derived piRNA (Josse et al. 2007). We initially tested the *BQ16* reporter, which is expressed predominantly in germline cells in the absence of piRNAs (Lemaitre et al. 1993; Josse et al. 2007). We evaluated *P{lacZ}* expression in both the earliest stages of oogenesis (contained within the germaria), and the developing egg chambers throughout the ovariole (Figure 3A). Silencing in the earliest stages of oogenesis is of particular interest because previous studies suggest that piRNA silencing for many germline TEs is transiently released in the germaria (3A; Dufourt et al. 2014). This reduction in silencing may pose a particular challenge to the regulation of *P-*elements, which actively transpose in the mitotic and pre-meiotic S-phase cells, which are contained within germaria (Spradling et al. 2011; Moon et al. 2018).

**Figure 3.**
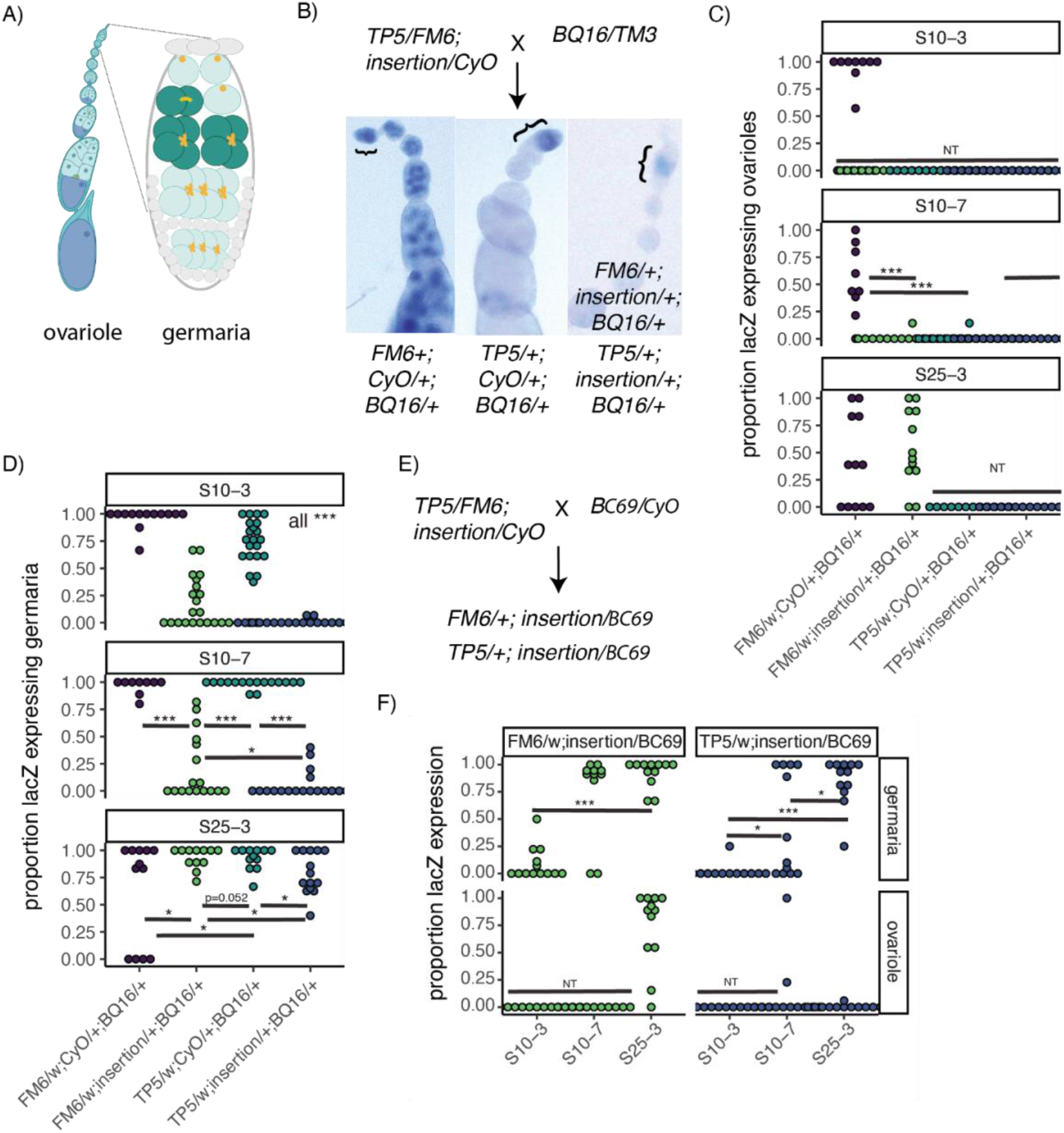
Silencing of *lacZ* reporters in germaria and ovarioles. A) Schematic of an ovariole and a germaria. Mitotically dividing cells in which *P*-elements transpose and piRNA silencing is reduced are highlighted in dark green. B) Crossing scheme to compare piRN A-like repression of the *BQ16lacZ* reporter between chromosomes containing *CR43651* insertions and the *TP5* chromosome containing a single P-element insertion in the *X-TAS* piRNA cluster. Representative images of different offspring genotype classes are also shown. Germaria, where piRNA silencing is incomplete and more robust in the presence of insertion chromosomes are indicated with brackets.The proportion of ovarioles (C) and germaria (D) with strong LacZ activity is compared between zygotic genotypes of the same mother. E) Crossing scheme to compare piRNA-like repression of the *BC69 lacZ* reporter among chromosomes containing *CR43651* with and without the *TP5* chromosome. F) The proportion of germaria and ovarioles with robust LacZ activity in the offspring from E. Genotypes were compared in panels C, D, and F through one-way permutation tests. NT indicated that the mean could not be compared to other groups due to absence of variance. Significant comparisons are indicated as follows: * P > 0.05, ** P > 0.01 *** P > 0.001. Panel A was created with BioRender.com

In the female germline, *trans* silencing of *P{lacZ}* reporters requires a repressive piRNA cluster insertion in both the maternal and zygotic genotype (Josse et al. 2008), allowing us to separate the zygotic effects of the insertion chromosomes from *TP5* by crossing *TP5/FM6; extraction chromosome/CyO* females to males with a *P{lacZ}* reporter (Figure 3B). Female offspring inheriting *TP5* but not extraction chromosomes (*w/TP5;+/CyO;BQ16/+*), exhibit no LacZ activity in developing egg chambers, but moderate LacZ activity in germaria (Figure 3C, 3D). This aligns with previous studies suggesting that telomeric *P-*elements silence the germline expression of *P{lacZ}* reporters, but that this silencing is incomplete in germaria (Josse et al. 2007; Dufourt et al. 2014). By contrast, female offspring that inherited neither *TP5* nor insertion chromosomes (*w/FM6; +/CyO; BQ16/+*) exhibited robust LacZ activity throughout the female germline.

Surprisingly, we observed that extraction chromosomes *S10-3* and *S10-7* also exhibit zygotic silencing effects, as flies that inherited these extraction chromosomes, but not *TP5* (*FM6/w; insertion chromosome/CyO*), exhibited strong repression of *P{lacZ}* (Figure 3B-D). Furthermore, *S10-3* and *S10-7* were associated with significantly stronger silencing in germaria than *TP5*, with the majority of germaria exhibiting weak or no LacZ activity (Figure 3B,D). By contrast, *S25-3* exhibited no silencing in germaria or ovarioles, as LacZ activity in these females did not differ from *w/FM6; +/CyO; BQ16/+* controls. Consistent with enhanced *P-*element piRNA production in the presence of all 3 insertion chromosomes (Figure 2), the most robust silencing is observed in flies inheriting both *TP5* and an insertion chromosome (*TP5/w; insertion chromosome/CyO*, Figure 3B,D).

To establish whether germaria silencing associated with *S10-3* and *S10-7* is general to germline expressed *P{lacZ}* reporters, we compared the ability of insertion chromosomes to silence the expression of a second reporter, *BC69. BC69* exhibits higher germline expression levels than *BQ16* and is less sensitive to piRNA-like silencing (Josse et al. 2008; Lemaitre et al. 1993). Because *BC69* is homozygous sterile and balanced by *CyO*, we could only compare silencing differences associated with the 3 insertion chromosomes in the presence and absence of *TP5* (Figure 3E,F). Similar to our results with *BQ16, S10-3* exhibited strong silencing in germaria regardless of whether TP5 was present or absent, whereas *S25-3* exhibited minimal silencing in germaria. *S10-7* exhibited an intermediate phenotype, in which germaria silencing did not differ from *S25-3* in the absence of *TP5*, however, silencing in germaria was stronger when *S10-7* was combined with *TP5*. These observations support our conclusion that *S10-3* and *S10-7*, but not *S25-3* confer *trans* silencing that is zygotically independent of *TP5*. They further suggest that *S10-3* may be a stronger silencer in germaria than *S10-7*.

### *P-element* transgenes in *CR43651* do not exhibit piRNA-like silencing

Our results suggest that chromosomes with antisense *P-*element insertions in *CR43651* show piRNA-like silencing that is reminiscent of—but more robust than—chromosomes carrying insertions into major piRNA clusters like *X-*TAS. Many large piRNA clusters, including TAS clusters, are characterized by a capacity to silence reporter genes *in trans* if DNA sequence homologous to the reporter is introduced into the piRNA cluster (Lemaitre et al. 1993; Josse et al. 2007; Muerdter et al. 2012). This property of TAS and other piRNA clusters highlights their function as a genetic trap for invading TEs. Silencing reflects the production of complementary piRNA from the cluster, which target homologous RNA for co-transcriptional and post-transcriptional silencing (Le Thomas, Stuwe, et al. 2014). For example, *P-*element derived transgenes located in TAS piRNA clusters are sufficient to establish silencing of *P{lacZ}* reporters in euchromatic regions (Josse et al. 2007; Ronsseray et al. 2001).

To determine if *CR43651* acts like a canonical piRNA cluster, regulating any sequences homologous to those inserted into this locus, we examined the capacity of 6 *P-*element derived transgenes into *CR43651* to silence *BQ16* in the ovary (F1 offspring of insertion females x *BQ16* males, Figure 4A, Supplementary Table 2). We observed that neither sense nor antisense *P-*element transgenic insertions in *CR43651* establish ovarian silencing of *BQ16* (Figure 4B-D). Since *P-*element derived transgenes and native *P*-elements exhibit equivalent sequence homologous to *P{lacZ}* (positions 84-586 of the native *P-*element), or more homologous sequence if the transgene also contains *lacZ*, these observations cannot be attributed to reduced homology with the transgenes themselves. Rather they suggest that antisense transcription from a *P-*promoter in *CR43651* alone is not sufficient to establish piRNA silencing of euchromatic targets like *BQ16*.

**Figure 4.**
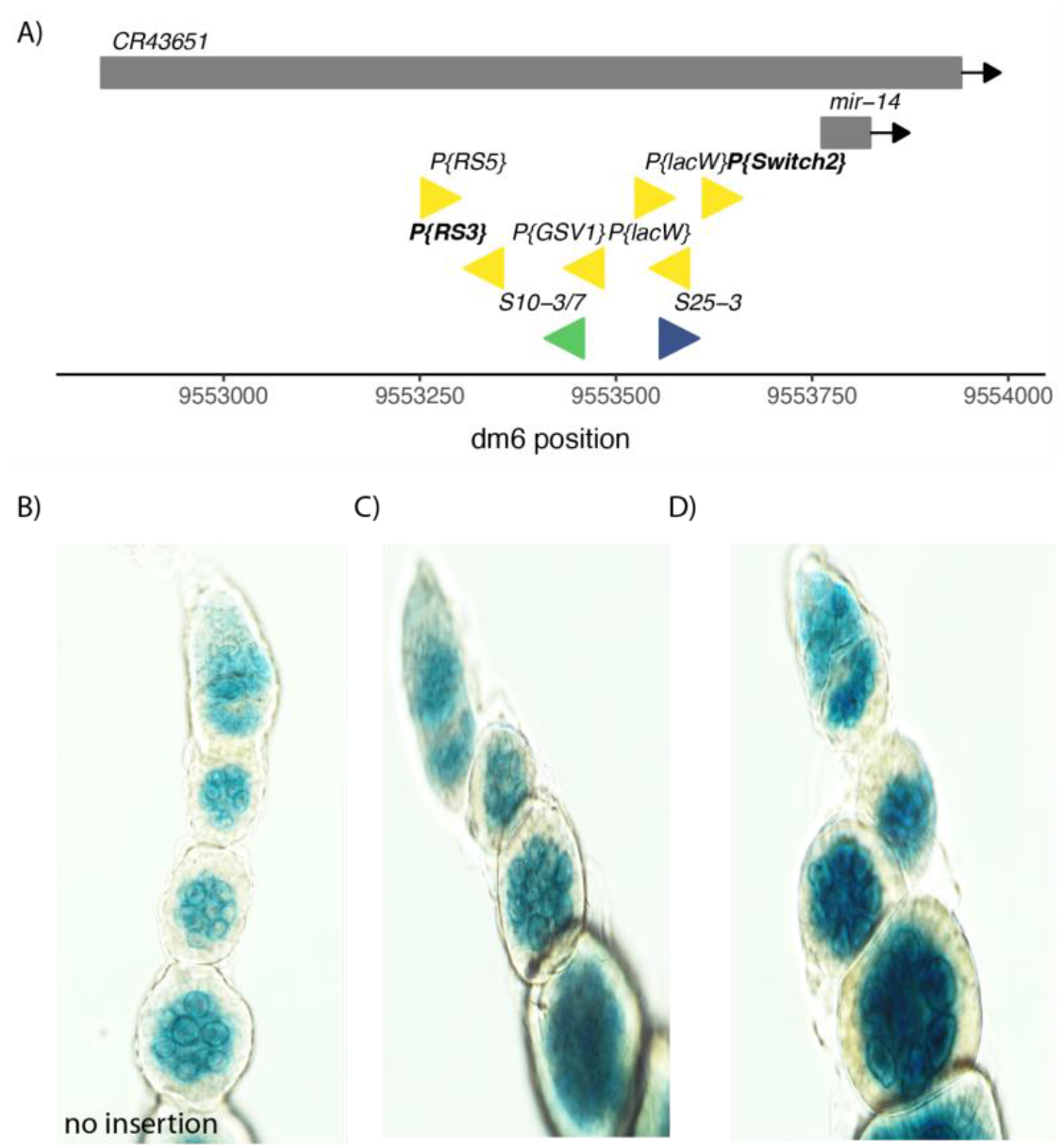
A) location and orientation of *P*-element insertions and transgenes within the *CR43651* locus. Insertions shown in C and D are indicated in bold. B-D) lacZ expression from the *BQ16* reporter in F1 ovaries from B) Oregon-RC (no insertion), C) *w[1118]*; *P{w[+mW*.*Scer\FRT*.*hs]=RS3}CB-5570-3*, and D) y*[1] w[∗]; P{w[+mW*.*hs]=Switch2}GSG3687-1/CyO* females crossed to *BQ16/TM3* males.

### Fitness costs associated with *CR43651* insertion chromosomes

In addition to establishing self-regulation through piRNA, TE insertions can also impact fitness, both positively and negatively, by altering the expression of adjacent host genes. The miRNA *miR14* is produced from *CR43651* precursor transcripts. To determine whether *P-* element insertions in *CR43651* reduce *miR14* expression, we examined miRNA abundance in *TP5* as compared to *TP5; insertion chromosome* homozygotes. We observed that for both biological replicates of all 3 insertion chromosomes, mature *miR14* (*miR14-RA*) is reduced ∼2-fold as compared to the *TP5* background (Figure 5A). While we cannot conclude that this reduced expression is due to *P-*element insertions, we observed only two other abundant miRNAs (mean RPKM across all samples >100, total 83 miRNAs) whose expression changed more than 1.5 in all 6 insertion chromosome samples (Supplementary Table 3). Thus, the magnitude and reproducibility of reduced *miR14-RA* expression is exceptional. Interestingly, the passenger strand miRNA (*miR14-RB*) was less affected in the insertion chromosomes, and did not change systematically from *TP5* abundance levels (Figure 5A). While this is puzzling given that both products are derived from the same pre-miRNA, passenger strand miRNAs are generally degraded (reviewed in Medley et al. 2021). Degradation is likely less regulated than the selection and retention of the active form, which might explain more variation *mir14-RB*.

**Figure 5.**
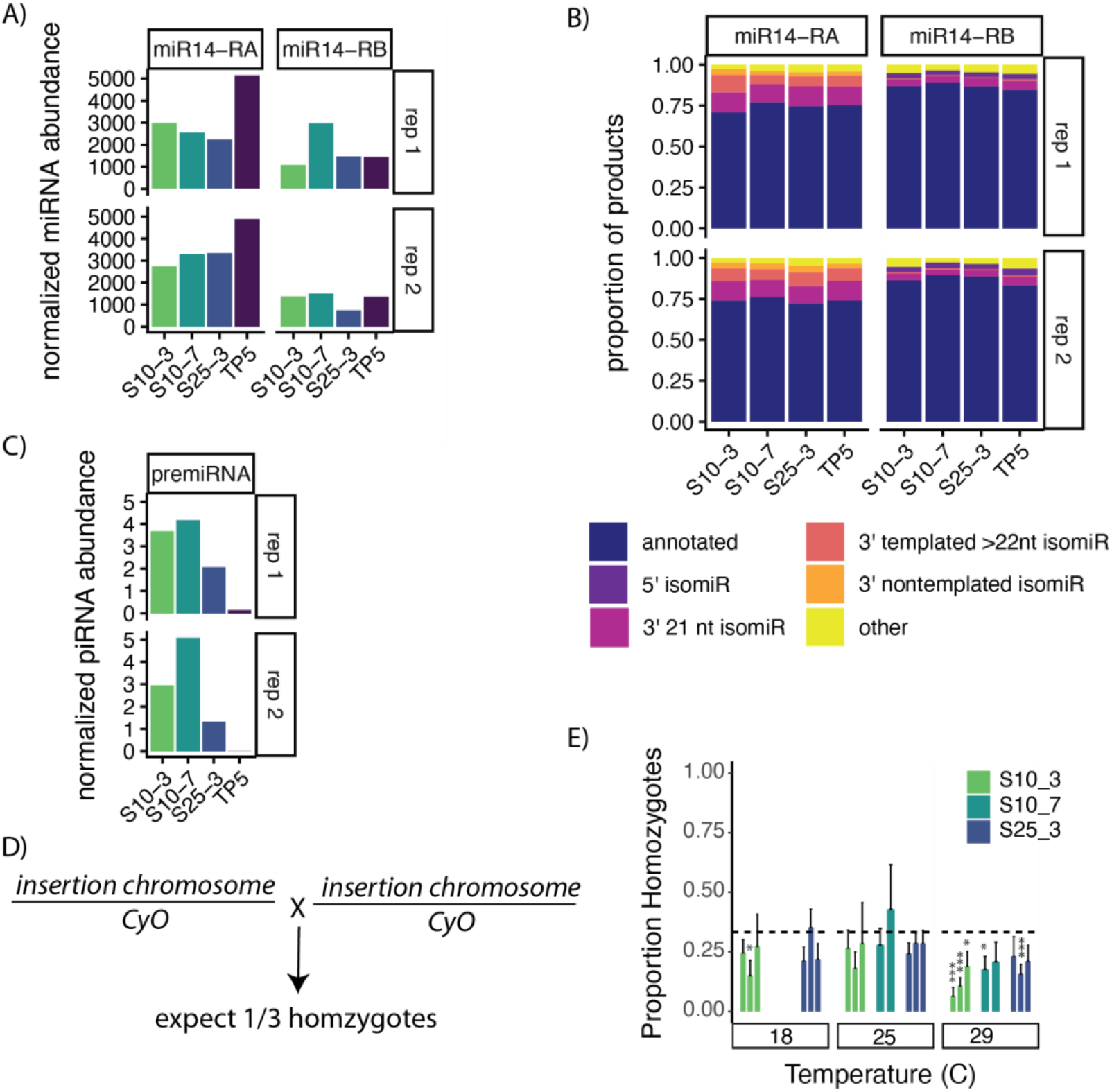
Chromosomes carrying insertions chromosomes show reduced *mirl4* expression and homozygous viability effects. A) Abundance and B) isomiR composition of mature *mir14* RNAs from insertion chromosomes as compared to the second chromosome in the *TP5* background. 5’ and 3’ isomiRs refer to a change in length relative to the annotated 22nt form, nontemplated isormRs include a terminal base that differs from that in the reference genome. C) Abundance of piRNAs overlapping the *mir14* pre-miRNA. Abundanceis A and C were normalized per million sequenced miRNAs. D) Crossing scheme to detect homozygous viability effects of insertion chromosomes. Assuming no effect on viability 1/3 of offpsring are expected to be homozygotes due to recessive lethality of the *CyO* balancer chromosome. E) Proportion homozygotes observed for each extraction chromosomes at 3 different temperatures. S1O_7 could only be replicated at higher temperatures since it was difficult to maintain the balancer chromosome in this stock. All insertion chromosomes are in a balanced *TP5* background *(TP5/FM6;insertion chromosome/CyO)*. P-values were obtained from a Chi-Squared Goodness-of-fit test given an expected probaility of 1/3. *,*** denotes P < 0.05, 0.001. small RNA sequencing underlying panel A-C are in PRJNA1292797.

*miR14-RA* levels could be reduced as a consequence of *P-*element insertions in *CR43651* in two ways: 1) reduced transcription as a consequence of piRNA mediated heterochromatin formation, or 2) aberrant processing of *miR14* pre-miRNAs into piRNAs. The latter scenario predicts that a significant fraction of products from *miR14* pre-miRNAs will correspond to piRNAs. We could not find evidence of this. In both *TP5* controls and insertion chromosomes, the overwhelming majority (∼95%) of aligned small RNAs (18-32nt) overlapping *miR14-RA* and *miR14-RB* correspond to the the annotated miRNA or an isomiR with an elongated, truncated or modified 5’ or 3’ end (Figure 5B). We furthermore found very few candidate piRNAs (>23nt and do not correspond to isomiRs) that overlap the *miR14* pre-miRNA, suggesting that the majority of this portion of the *CR43651* transcript is correctly processed into miRNAs (Figure 5C).

Nevertheless, rare piRNAs do arise from the *miR14* pre-miRNA, and these piRNA are increased with insertion chromosomes (Figure 5C). *mir14-*derived piRNA are overwhelmingly in the sense orientation: only two antisense piRNAs overlapping the *miR14* pre-miRNA were sequenced across all samples and replicates. This is consistent with the establishment of piRNA production from this locus by *P-*element insertions, as piRNA biogenesis spreading from stand alone TEs is generally unistranded, arising from piRNA transcription originating in the TE insertion (Olovnikov et al. 2013; Shpiz et al. 2014).Thus, it does not appear that *mir14* is a direct target of antisense piRNA. Rather, our data suggest that a small minority of sense *CR43651* transcript is processed in piRNA through phased piRNA biogenesis (Mohn et al. 2015), initiated by upstream *P-*element insertions. Reduced abundance of *mir14-RA* therefore likely arises through heterochromatin formation, which is coincident with *de novo* piRNA biogenesis (Olovnikov et al. 2013; Le Thomas, Stuwe, et al. 2014; Le Thomas et al. 2013; Shpiz et al. 2014; Mohn et al. 2014).

*miR14* has a number of known functions in cell death, metabolism and development, particularly in larval stages (Varghese & Cohen 2007; Xu et al. 2003; Nelson et al. 2014). As a result, *miR14* mutants furthermore generally die at the pupal stage (Xu et al. 2003). To determine whether homozygosity for *P-*element insertions into *CR43651* could impact egg to adult viability, we compared the eclosion rates of insertion chromosome homozygotes (*TP5;insertion chromosome*) to their heterozygous siblings (*TP5;insertion chromosome/CyO*) at 3 different temperatures (18°C,22°C,29°C). All insertions have significant homozygous fitness costs at 29°C, with fewer than the expected ⅓ of the eclosing adults corresponding to homozygotes (⅓ is expected because *CyO* homozygotes are not viable, Figure 5D,E). One replicate of *S10-3* also exhibited fitness costs at 18°C. While we cannot rule out that these fitness effects are impacted or exacerbated by homozygosity for other *P-*element insertions on these same chromosomes, the only insertion site shared between all three chromosomes is *CR43651*. This strongly suggests that homozygosity for *P-*element insertions in *CR43651* has deleterious effects on egg to adult viability, consistent with the reduction in *miR14* expression we observe. Consistent with a partial loss of *mir14* function, these viability effects are mild: on average across all samples there was 50% lethality at 29°C and only 27% at 18°C. In contrast, homozygous *miR14* mutants exhibit 80% lethality (Xu et al. 2003).

## DISCUSSION

Our previous work demonstrated that *P-*element insertions into the long-noncoding RNA *CR43651* represent beneficial mutations in laboratory populations that have recently been invaded by *P-*elements (Wang et al. 2023). In this work we explore the phenotype associated with *CR43651* insertion alleles and uncover unique silencing properties of these chromosomes. Our work reveals that *CR43651* is not a piRNA cluster in the traditional sense, adds to a growing body of literature that blurs the distinctions between piRNA clusters and piRNA targets (Olovnikov et al. 2013; Gebert et al. 2021; Srivastav et al. 2024), and support a recently proposed model for how new piRNA clusters emerge (Srivastav et al. 2024).

### Why are *P-*element insertions in *CR43651* beneficial?

We previously showed that *P-*element insertions in *CR43651* are likely beneficial in laboratory populations, owing to the unusually large number of insertion alleles at this locus, as well as their high population frequencies (Wang et al. 2023). Our findings here suggest that one possible benefit of *P-*element insertion alleles in *CR43651* is their unusual *trans* silencing properties in the germaria. Germaria house the S-phase cells where *P-*elements transpose (Spradling et al. 2011; Moon et al. 2018), making them profoundly significant for *P-*element control. Furthermore, piRNA silencing established by insertions in canonical TAS piRNA clusters, like the *TP5* insertion, is incomplete in germaria (Figure 3; Dufourt et al. 2014). *CR43651* insertion chromosomes therefore have obvious potential benefits for the control of *P-* elements and other DNA transposons that mobilize during S-phase.

Alternatively, the adaptive advantage of *P-*element insertions in *CR43651* could arise from the altered expression of *miR14*, which is produced from the *CR43651* precursor. *miR14*, which is involved in regulating numerous developmental and life history traits (Xu et al. 2003; Varghese & Cohen 2007; Nelson et al. 2014). Our data suggest that insertion alleles into *CR43651* reduce the production of the guide strand *miR14-RA* (Figure 5A). However, our data do not suggest that these effects are beneficial. All three chromosomes containing *P*-element insertions into *CR43651* exhibited homozygous viability effects (Figure 5D,E). We cannot conclusively attribute these viability reductions to reduced production of *miR14*. However they are consistent with *miR14* loss of function, as *miR14* mutants show a high rate of pupal lethality (Xu et al. 2003; Varghese & Cohen 2007; Nelson et al. 2014).

Collectively, our data suggest that *P-*element insertion alleles in *CR43651* are subject to antagonistic pleiotropy: conferring dominant benefits in *P-*element repression, and subject to negative selection in homozygotes due to effects on *mir14-RA*. While this model is purely speculative, it would explain why *P-*element insertion alleles in *CR43651* are so common in our evolving laboratory populations, but are comparatively rare in recent wild-derived inbred lines. Of >200 inbred lines in the *Drosophila* Genetic Reference Panel (DGRP), only one strain contains a *P-*element insertion allele in *CR43651* (Mackay et al. 2012; Zhang et al. 2020). Fitness costs of homozygosity for these insertions may explain why this is so.

### Do some *P-*element insertion alleles in *CR43651* represent piRNA clusters?

*CR43651* is not considered a piRNA cluster, as this locus is not generally a source of piRNA, including in the A4 strain in which we performed our laboratory evolution experiments (Wang et al. 2023; Srivastav et al. 2024). However, our work here establishes that *P-*element insertions alleles of *CR43651* can exhibit two major features of piRNA clusters: 1) their ability to produce piRNA and 2) their ability to target silencing in *trans*. piRNA clusters are extraordinarily polymorphic within populations, with the same genomic regions acting as a source of piRNAs in some strains but not others (Srivastav et al. 2024; Signor et al. 2023; Wierzbicki et al. 2023; Gebert et al. 2021; Rozhkov et al. 2010; Erwin et al. 2015). We propose that the antisense insertion allele in *CR43651* that we investigated here may represent a piRNA cluster in its infancy, providing insights into how those clusters may emerge.

With regards to piRNA biogenesis, we showed that in a *TP5* background, *CR43651* insertions exhibit divergent piRNA production adjacent to the insertion site. This signature of piRNA production spreading from a euchromatic TE into adjacent genomic sequence is characteristic of *de novo* piRNA cluster formation (Shpiz et al. 2014; Mohn et al. 2014). It is intriguing that this feature was not general to other *P-*element insertions on the same chromosome, rather it is an unusual property of *CR43651*. One obvious explanation for this is the regulatory environment: *CR43651* may contain enhancers that are conducive to *P-*element expression and piRNA production. However, the orientation of the insertion may also matter, since we observe that chromosomes harboring the antisense insertion in *CR43651 (S10-3* and *S10-7*) exhibited stronger divergent piRNA production (Figure 1B), and more robust amplification of *P-*element derived piRNAs (Figure 2A-C) than the chromosome with a sense insertion (*S25-3*). Convergent transcription leads to the production of dsRNA, is known to be important for the regulation of some germline piRNA clusters (Andersen et al. 2017), and may play a role in the formation of new piRNA clusters (Luo et al. 2023).

Consistent with their piRNA production, our reporter gene assays reveal that chromosomes harboring antisense *P-*element insertions in *CR43651* exhibit a second major feature of piRNA clusters: *trans-*silencing properties. While we cannot attribute this silencing exclusively to antisense *CR43651 P-*element insertions, other insertions found on the same chromosomes do not appear to act as source loci for piRNA (Figure 2A,C, Supplementary Figure 1). Critically, *trans-*silencing exhibited by antisense insertion chromosomes does not require zygotic *TP5*, suggesting that *CR43651* insertions have silencing properties that are at least partially separable from an insertion in a major piRNA cluster (Figure 3B-E). However, unlike major piRNA clusters such as *flamenco* and *X-TAS*, the transgenic insertion of exogenous sequences alone is not sufficient to establish silencing of a reporter in *trans* (Figure 4). Hence, unlike *TAS* clusters, *CR43651* is not a trap. This is not surprising because *CR43651* is not a source of piRNAs in the absence of *P-*element insertions (Figure 1B).

We speculate that exposure to the *P-*element piRNA from trap insertions may have facilitated the conversion of *CR43651* into a piRNA cluster. Maternally transmitted piRNAs are known to establish homology dependent epigenetic changes at both non-allelic and allelic loci, allowing them to produce piRNA and target silencing *in trans* (de Vanssay et al. 2012; Hermant et al. 2015; Dorador et al. 2022; Le Thomas, Tóth, et al. 2014; Erwin et al. 2015). Srivastav et al.(2024) recently proposed a model for the emergence of new piRNA clusters. Following the introduction of TE copies into major piRNA cluster “traps”, new piRNA producing loci emerge around the genome through the epigenetic conversion of some new target TEs into piRNA clusters. Over time, these clusters may incorporate additional TE insertions, grow in size and become more stable allowing them to function independently of the original trap insertion (Srivastav et al. 2024; Gebert et al. 2021). In the experimental populations where we discovered these *CR43651 P-*element insertions, insertions into *TAS* “trap” piRNA clusters were also very common (Wang et al. 2023). *P-*element insertions in *CR43651* may therefore represent the first step in piRNA cluster formation: the production of a new source locus with *trans* silencing ability.

## MATERIALS AND METHODS

### Generation of chromosome extraction lines and quantification of viability effects

To isolate individual second chromosomes containing *P-*element insertions in *CR43651*, we extracted them using *w[1118]; wg[Sp-1]/CyO; sens[Ly-1]/TM6B, Tb[1]* (S2 Table). We then used a *TP5:WgSp-1/CyO* strains to move second chromosomes into a *TP5* background. We then screened for individual chromosome extraction lines containing insertions into *CR43651* using two PCR primers that flank the insertion hotspot region to look for insertion lines with unexpectedly large amplicons (Wang et al. 2023). We then characterized whether the insertions were on the plus or minus strand by combining primers in the *CR43651* with primers in *P-* element (Supplementary Table 4). Stocks were maintained as *TP5;P{}CR43651/CyO*. To estimate viability effects, flies were intercrossed in temperature controlled incubators, and the number of heterozygous *TP5;P{}CR43651/CyO* and homozygous *TP5;P{}CR43651* offspring were quantified.

### *PLacZ* silencing assays

*BC69* and *BQ16* are *P{lacZ}* reporter genes on the second and third chromosomes that are expressed in the female germline. They contain in-frame fusions between the second exon of *P-transposase* and *Escheria coli lacZ* (O’Kane & Gehring 1987), rendering their LacZ production sensitive to the presence of piRNA (Josse et al. 2007). *TP5;P{}CR43651/CyO* females were crossed to *TP5;wg*^*Sp-1*^*/CyO* males to isolate double-balanced females: *TP5/FM6;wg*^*Sp-1*^*/CyO*. These females were then crossed to *BC69/CyO* and *BQ16* males, and F1 females were collected and reared on yeast paste for 2-4 days. For crosses with *w;BQ16/Sb* compared all possible combination of first and second chromosomes, but for *w;BC69/CyO*, we compared only *TP5/w;P{}CR43651/BC69* with *FM6/w;P{}CR43651/BC69*, because we could not differentiate between *P{}CR43651/CyO* and *BC69* genotypes.

To ascertain silencing from *P-*element derived transgenes in *CR43651*, we crossed homozygous and balanced transgenic stocks (S2 Table) to *w;BQ16/Sb* males only and examined LacZ activity in offspring containing both the *CR43651* transgene and *w;BQ16/Sb*.

Dissected ovaries were stained for LacZ activity as in Lemaitre *et al*. (1993). Briefly, ovaries were dissected in 1X PBS and fixed in 0.5% glutaraldehyde for 4-6 minutes. They were then washed in 1X PBS twice, then incubated in 0.2% X-Gal overnight and 37°C. Ovaries were then washed and mounted in glycerol. Stained ovaries were visualized on a Nikon SZX10 under 20X magnification, or on a Nikon Ti2 under 60X magnification. LacZ expression was scored separately for germaria and developing egg chambers. For statistical analysis of LacZ activity, the proportion of LacZ expressing germaria or ovarioles was compared between genotypes, based on ∼9 ovarioles from ∼16 females for each genotype for BQ16 crosses, and based on ∼13 ovarioles from ∼12 females for BC69. The proportion of lacZ expressing ovarioles was compared between genotypes using a one-way permutation test implemented in the R-package “coin”.

### Small RNA sequencing and analysis

Two biological replicates including ovaries from 20 3-5 day old females maintained with yeast paste were dissected for each of the extraction chromosomes in the *TP5* background (*TP5;P{}CR43651*), as well as in the *TP5* background without the transgene. Ovaries were homogenized in Trizol and RNA was extracted according to manufacturer instructions, except that RNA was precipitated overnight with an excess of ice-cold ethanol. Depletion of 2S rRNA was completed according to (Wickersheim & Blumenstiel 2013), and small RNA library prep and single-end 50 nt sequencing on the Novaseq X plus was completed by Novogene (Carlsbad, CA).

Adapters were removed from small RNA sequencing reads, and they were separated into 2 size classes with trim-galore, 18-22 nt: and 23-32 nt. Read alignments were performed using BWA (Li & Durbin 2009). We first removed ribosomal RNAs (rRNAs) by aligning and removing reads that aligned to any annotated rRNA, including 2sRNA, from Release 6 of the *D. melanogaster* genome (dm6) (Hoskins et al. 2015). miRNAs were then identified based on their alignment to all annotated miRNAs in dm6 (up to 1 mismatch). 23-32 nt RNAs did not correspond to miRNAs but aligned to the A4 assembly, a repbase TE consensus sequence for *D. melanogaster* (Bao et al. 2015), or the *P-*element consensus (0 mismatches), were considered piRNAs. For sliding window analyses and *P*-element derived piRNA abundance, raw read counts were normalized as reads per million mapped miRNAs from the same library.

Sliding window analysis of piRNA abundance over *P*-elements or their insertion sites were calculated using bedtools genomecov (Quinlan & Hall 2010). Ping-pong abundance was identified using an in-house script from Kelleher *et al* (2012).

### Genome Sequencing and Identification of *P-*element insertion sites

Illumina genomic libraries were generated with the Nextera CD Index kit (Illumina, San Diego, CA) according to manufacturer instructions. 150 nt Paired-End Sequencing was performed on a Next-Seq 500 by the University of Houston Seq ‘n Edit core. Adapters were removed using trim-galore. *P*-element insertions in A4 were identified, and their frequencies were estimated within each strainy through the PIDFE pipeline (Zhang & Kelleher 2017; Zhang et al. 2020; Wang et al. 2023), and are reported in Supplementary Table 1.

## ACKNOWLEDGEMENTS

We are grateful to Stéphane Ronsseray and Antoine Boivin for providing the *P{lacZ}* reporter strains *BQ16* and *BC69*. This work was supported by R35 GM138112 to ESK. NMB and MW were supported by NSF 2147073 to Richard Meisel.

