## Supplementary Material for "Adaptive *P*-element insertions in a long non-coding RNA are an emerging piRNA cluster"

| A4 position | strand | chromosome | freq | dm6 chromos | dm6 coordinate | piC Srivastav |
|---|---|---|---|---|---|---|
| 5038438 | - | S25-3 | 1 | 2L | 5027479 | no |
| 9792318 | + | S25-3 | 1 | 2L | 9782275 | 1 replicate onl |
| 13399471 | + | S10-7 | 1 | 2L | 13399017 | no |
| 15064056 | + | S25-3 | 1 | 2L | 15074469 | no |
| 15064137 | - | S25-3 | 1 | 2L | 15074550 | no |
| 15663748 | + | S10-3 | 0.37 | 2L | 15731853 | no |
| 18937845 | - | S25-3 | 0.92 | 2L | 18943951 | no |
| 1010919 | - | S10-3/S10-7 | 1/1 | 2R | 1210315 | no |
| 6505507 | + | S10-3/S10-7 | 1/0.96 | 2R | 6740685 | no |
| 9279761 | - | S10-3/S10-7 | 0.96/0.95 | 2R | 9553459 | no |
| 9279848 | + | S25-3 | 0.97 | 2R | 9553556 | no |
| 12927939 | - | S25-3 | 1 | 2R | 13225050 | no |
| 17120926 | + | S10-3 | 0.96 | 2R | 17394577 | no |
| 18331072 | + | S25-3 | 1 | 2R | 18611846 | no |
| 18480221 | - | S10-7 | 1 | 2R | 18761125 | no |
| 23308418 | - | S10-3 | 0.16 | 2R | 23548270 | no |
| 24761327 | - | S10-7 | 0.99 | 2R | 25032351 | no |
| 21750356 | + | S10-3 | 0.11 | 3L | 21925540 | no |

**Supplementary Table 1**. Location of P-element insertions on extraction chromosomes in A4 and dm6, and whether these coordinates occur in a known piRNA cluster in A4.

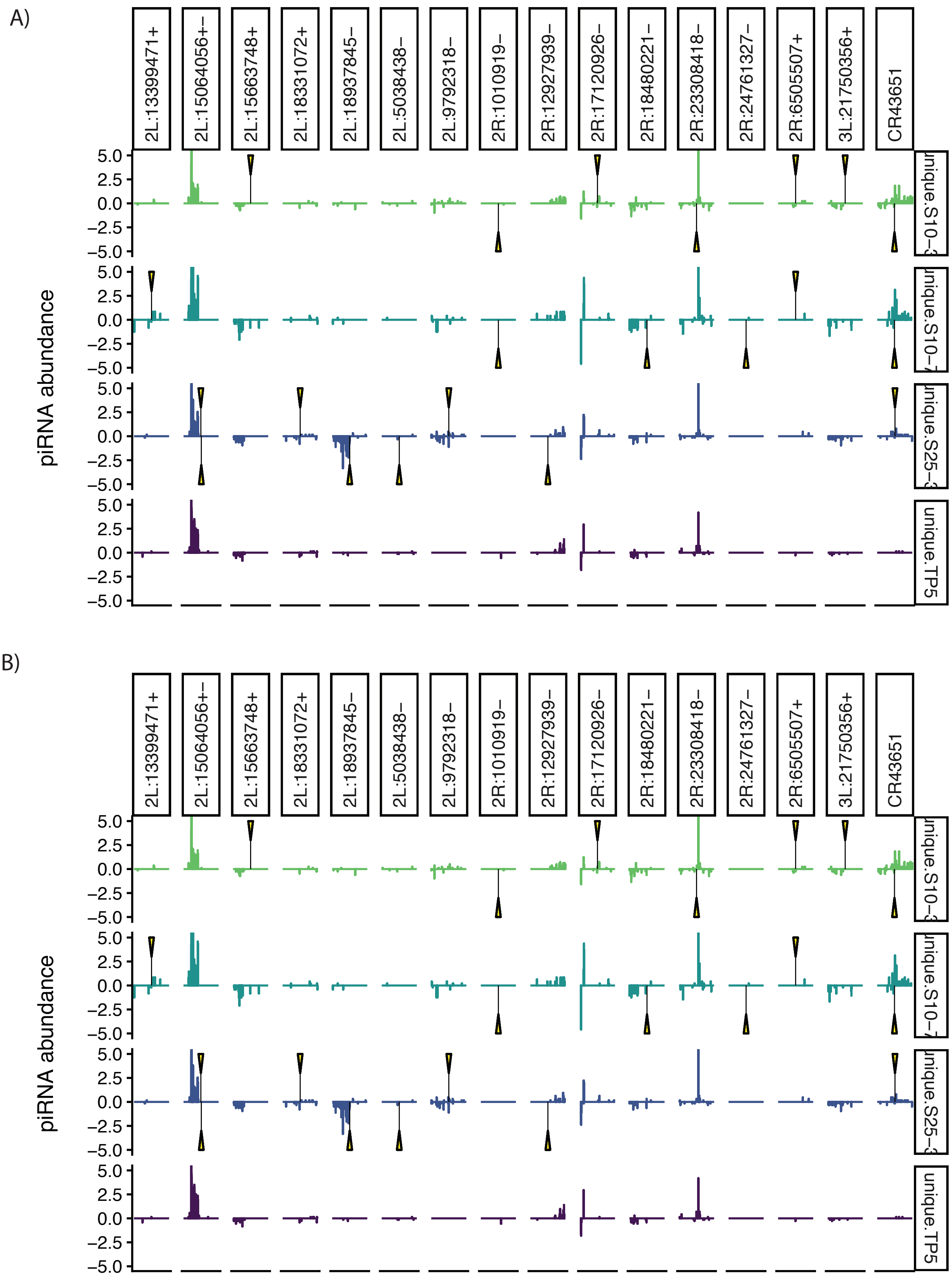


**Supplementary FIgure 1.** Sliding window analysis of uniquely mapping piRNAs aligning to the 6 Kb window surrounding insertion site. Panels A and B show separate biological replicates. Genome and small RNA sequencing underlying all panels are in PRJNA1292797.

| strain type | genotype | source | stock number |
|---|---|---|---|
| CR43651 transgenic insertion | *w[1118]; P{w[+mW.Scer\FRT.hs]=RS3}CB-5570-3* | kyoto | 123-691 |
| CR43651 transgenic insertion | *w[1118]; P{w[+mW.Scer\FRT.hs]=RS5}5-HA-1040* | kyoto | 125-025 |
| CR43651 transgenic insertion | *y[1] w[67c23]; P{w[+mC]=lacW}lncRNA:CR43651[k01301]/CyO* | BDSC | 10502 |
| CR43651 transgenic insertion | *w[*]; P{w[+mC]=lacW}mir-14[k10213]/CyO, P{ry[+t7.2]=sevRas1.V12}FK1* | BDSC | 10982 |
| CR43651 transgenic insertion | *y[1] w[*]; P{w[+mW.hs]=Switch2}GSG3687-1/CyO* | BDSC | 40327 |
| CR43651 transgenic insertion | *w[*]; P{w[+mGS]=GSV1}lncRNA:CR43651[EP-65]* | BDSC | 43435 |
| *P{lacZ}* reporter | *BQ16/Sb+* | Stephane Ronsseray | NA |
| *P{lacZ}* reporter | *BC69* | Antoine Boivin | NA |
| balancer | w[1118]; wg[Sp-1]/CyO; sens[Ly-1]/TM6B, Tb[1] | BDSC | 33821 |
| P-element piRNA regulation | *TP5* | BDSC | 64168 |
| P-element piRNA regulation | *TP5; wg[Sp-1]/CyO* | BDSC | 64168 |

**Supplementary Table 2.** strains used in this study.

| FBtr | S10.3.3FC | S10.3.4FC | S10.7.3FC | S10.7.4FC | S25.3.3FC | S25.3.4FC | grand_mean |
|---|---|---|---|---|---|---|---|
| FBtr03042 | 0.1 | 0.02 | -0.46 | -0.22 | -0.24 | 0 | 180579.2 |
| FBtr03043 | -0.34 | 0.2 | 0.25 | -0.03 | 0.65 | -0.47 | 146244.7 |
| FBtr03044 | 0.05 | 0.09 | -0.42 | -0.09 | 0.04 | -0.01 | 121043.7 |
| FBtr03042 | -0.45 | -0.49 | -0.92 | -0.71 | -1.29 | -0.16 | 49396.29 |
| FBtr03044 | -0.12 | 0.08 | 0.57 | -0.07 | 0.22 | -0.49 | 48860.89 |
| FBtr03043 | -0.06 | 0.03 | 0.96 | 0.02 | 0.25 | -0.63 | 47474.03 |
| FBtr03044 | 0.22 | 0.33 | 0.87 | 0.1 | 0.29 | -0.46 | 41914.47 |
| FBtr03042 | 0.22 | 0.29 | 0.73 | 0.16 | -0.08 | -0.3 | 41185.16 |
| FBtr03043 | -0.23 | -0.02 | -0.19 | -0.04 | 0.27 | -0.29 | 36901.69 |
| FBtr03042 | 0 | -0.09 | -0.13 | 0.42 | 0.25 | 0.4 | 34169.57 |
| FBtr03043 | 0.37 | 0.39 | 0.8 | 0.17 | 0.28 | -0.28 | 32311.31 |
| FBtr03043 | 0.03 | -0.59 | -0.16 | 0.83 | -0.62 | 0.48 | 29319.2 |
| FBtr03042 | 0.07 | -0.24 | -0.84 | -0.07 | -0.45 | 0.45 | 21929.29 |
| FBtr03044 | 0.82 | -0.93 | -3.54 | 0.69 | -0.71 | 1.54 | 16684.51 |
| FBtr03043 | -0.03 | -0.41 | 0 | 0.35 | -1.3 | 0.71 | 15754.95 |
| FBtr03044 | -0.03 | 0.39 | 1.32 | 0.34 | 0.44 | -0.5 | 15703.71 |
| FBtr03044 | 0.12 | -0.64 | -0.36 | 0.4 | -0.77 | 0.61 | 14955.56 |
| FBtr03043 | 0.07 | 0.02 | -0.21 | 0.09 | 0.3 | 0.43 | 8746.19 |
| FBtr03041 | 0.32 | -0.42 | -0.32 | 0.49 | -0.73 | 0.68 | 8348.7 |
| FBtr03042 | 0.34 | 0.4 | -0.53 | -0.07 | 0.14 | 0.28 | 6515.72 |
| FBtr03044 | 0.34 | 0.39 | -0.59 | -0.05 | 0.14 | 0.26 | 6489.36 |
| FBtr03044 | -0.11 | -0.7 | -0.61 | -0.17 | -0.95 | 0.52 | 5636.53 |
| FBtr03045 | 0.37 | -0.51 | -0.39 | 0.59 | -0.74 | 0.84 | 5245.07 |
| FBtr03044 | -0.05 | -0.52 | -0.7 | -0.41 | -0.39 | 0.09 | 5095.81 |
| FBtr03043 | 0.52 | 0.29 | -2.01 | 0.51 | -0.73 | 1.42 | 4183.18 |
| FBtr03044 | -0.78 | -0.93 | -1.03 | -0.61 | -1.23 | -0.59 | 3811.81 |
| FBtr03044 | -0.2 | -0.69 | -0.74 | -0.27 | -0.51 | 0.24 | 3505.83 |
| FBtr03044 | -0.08 | -0.63 | -0.78 | 0.06 | -0.66 | 0.37 | 3466.74 |
| FBtr04727 | -0.02 | 0.01 | -0.48 | -0.1 | -0.09 | -0.06 | 3378.4 |
| FBtr03045 | 0.44 | 0.1 | -0.51 | 0.02 | -0.32 | 0.48 | 3141.15 |
| FBtr03043 | -0.1 | -0.71 | -1.09 | -0.13 | -1 | 0.47 | 2422.15 |
| FBtr04727 | 0.11 | 0.24 | 0.4 | 0.13 | -0.01 | -0.15 | 1813.85 |
| FBtr04727 | -0.35 | -0.07 | 1.02 | 0.13 | 0 | -0.87 | 1733.35 |
| FBtr04727 | -0.17 | -0.15 | 0.23 | 0 | 0.05 | -0.69 | 1686.43 |
| FBtr03045 | -1.14 | -1.1 | -1.06 | -1.21 | -0.9 | 0.15 | 1646.45 |
| FBtr03044 | -0.1 | -0.68 | -1.09 | -0.42 | -1.04 | 0.7 | 1600.99 |
| FBtr03043 | -0.09 | -0.62 | -0.88 | -0.5 | -0.73 | 0.3 | 1593.88 |
| FBtr03045 | 0.53 | 0.26 | -0.79 | 0.16 | -0.16 | 0.53 | 1501.61 |
| FBtr03042 | 0.53 | 0.23 | -0.81 | 0.18 | -0.14 | 0.52 | 1496.64 |
| FBtr03042 | 0.31 | 0.13 | -0.84 | -0.13 | -0.23 | 0.48 | 1320.51 |
| FBtr03041 | 0.57 | -0.32 | -0.61 | 0.77 | -0.54 | 0.97 | 1310.21 |
| FBtr03043 | 0.59 | -0.3 | -0.62 | 0.73 | -0.52 | 0.98 | 1305.02 |
| FBtr03044 | -1.9 | -2.42 | -2.92 | -2.3 | -2.66 | -1.59 | 1085.2 |
| FBtr03041 | 0.6 | 0.01 | -0.33 | 0.31 | -0.27 | 0.91 | 1060.88 |
| FBtr03044 | 0.66 | 0.32 | 0.52 | 0.23 | 0.17 | 0.26 | 1028.44 |
| FBtr03042 | 0.33 | 0.03 | -0.65 | 0.01 | -0.11 | 0.42 | 1021.4 |

| | | | | | | | |
|---|---|---|---|---|---|---|---|
| FBtr030431 | -0.27 | -0.32 | -0.57 | 0.02 | -0.03 | 0.13 | 960.6 |
| FBtr047272 | 0.55 | 0.44 | -0.37 | 0.5 | 0.08 | 0.48 | 872.75 |
| FBtr030422 | 0.38 | 0.44 | -0.61 | -0.05 | -0.15 | 0.23 | 746.28 |
| FBtr030438 | 0.21 | 0.01 | -2.17 | 0.72 | -1.09 | 1.31 | 662.51 |
| FBtr047272 | 0.18 | 0.03 | -0.65 | 0.16 | -0.14 | 0.51 | 494.32 |
| FBtr030428 | 0.13 | -0.47 | -0.33 | 0.37 | -0.21 | 0.71 | 465.13 |
| FBtr047275 | -0.15 | 0.04 | 0.29 | 0.04 | 0.04 | -0.55 | 426.4 |
| FBtr047271 | -0.07 | 0.47 | 1.71 | 0.42 | 0.36 | -0.58 | 424.28 |
| FBtr047275 | 0.4 | 0.41 | 1.68 | 0.74 | 0.39 | -0.26 | 420.3 |
| FBtr030449 | -0.3 | -0.16 | 0.11 | -0.21 | -0.02 | -0.15 | 418.76 |
| FBtr047271 | -0.29 | 0.02 | -0.35 | -0.22 | 0.08 | -0.03 | 407.37 |
| FBtr047275 | 0.19 | 0.14 | -0.29 | -0.22 | 0.02 | 0.08 | 390.17 |
| FBtr047281 | -0.02 | -0.32 | 0.66 | 0.94 | 0.04 | 0.11 | 370.32 |
| FBtr030428 | 0.11 | -0.42 | -0.79 | 0.46 | -0.66 | 0.62 | 369.44 |
| FBtr030436 | -0.03 | -0.2 | -0.67 | 0.17 | -0.2 | 0.19 | 349.32 |
| FBtr030424 | -0.2 | 0.14 | 0.34 | -0.02 | 0.25 | -0.27 | 328.21 |
| FBtr030428 | 0.23 | -0.51 | -0.61 | 0.28 | 0.11 | 0.64 | 288.6 |
| FBtr030437 | 0.19 | -0.56 | -0.68 | 0.22 | 0.03 | 0.68 | 285.87 |
| FBtr030444 | -0.12 | 0.36 | 0.57 | -0.07 | 0.18 | -0.71 | 264.26 |
| FBtr030435 | 0.3 | 0.42 | -0.72 | 0.1 | 0.13 | 0.52 | 261.97 |
| FBtr047275 | 0.26 | 0.02 | 0.48 | 0.37 | 0.08 | -0.01 | 259.69 |
| FBtr030418 | 0.66 | -1.1 | -2.12 | 1.36 | -0.96 | 1.84 | 258.33 |
| FBtr030974 | 0.23 | -0.39 | -0.45 | 0.45 | -0.66 | 0.58 | 246.77 |
| FBtr030419 | -0.78 | -0.89 | -0.45 | 0.79 | -1.05 | 1.15 | 218.17 |
| FBtr030437 | 0.46 | -0.2 | -0.62 | -0.04 | 0.21 | 0.57 | 192.94 |
| FBtr047280 | -0.19 | -0.58 | -0.21 | 0.23 | -0.83 | 0.57 | 180.62 |
| FBtr030425 | 0.16 | -0.64 | -0.25 | 0.9 | -0.57 | 0.52 | 176.42 |
| FBtr030433 | 0.8 | -1.15 | -5.01 | 1.37 | -1.9 | 2.28 | 174.51 |
| FBtr047271 | -0.03 | -0.05 | 0.31 | -0.29 | -0.39 | -0.24 | 171.97 |
| FBtr030420 | 0.2 | 0.09 | 0.6 | 0.09 | 0.32 | -0.32 | 156.98 |
| FBtr030432 | 1.41 | 0.98 | 1.27 | 1.5 | 1.39 | 1.23 | 155.36 |
| FBtr030422 | 0.01 | 0.18 | -0.58 | -0.05 | -0.15 | 0.14 | 140.96 |
| FBtr047274 | 0.31 | -0.55 | -0.12 | 0.41 | -0.35 | 0.43 | 140.53 |
| FBtr047273 | -0.19 | -0.32 | -0.43 | 0.42 | -0.47 | 0.17 | 125.35 |
| FBtr030432 | -0.07 | -0.1 | -0.59 | -0.12 | 0.1 | -0.09 | 122.71 |
| FBtr047272 | 0.73 | 0.57 | 1.5 | 1.19 | 0.52 | -0.05 | 113.19 |
| FBtr047275 | 0.08 | 0.21 | 1.21 | 0.57 | 0.48 | -0.37 | 101.46 |

Supplementary Table 3. FC in expression of 83 highly expressed miRNAs in each of 4 genotypes.

| Insertion | Location | Forward Primer Name | Target | Forward Primer Sequence | Reverse Primer Name | Reverse Primer Target |
|---|---|---|---|---|---|---|
| sense | 5' | 2RlncRNACR43651-F1 | CR43651 | 5'-GAGAGAGAGAGCGCGTCCTA-3' | P-element5-R2 | P-element |
| sense | 3' | P-elementI-F1 | P-element | 5'-GAAAATGCCACCGAAACTGC-3' | 2RlncRNACR43651-R1 | CR43651 |
| antisense | 5' | P-element5-R2 | P-element | 5'-TTGGGAGTTTTCACCAAGGC-3' | 2RlncRNACR43651-R1 | CR43651 |
| antisense | 3' | 2RlncRNACR43651-F1 | CR43651 | 5'-GAGAGAGAGAGCGCGTCCTA-3' | P-elementI-F1 | P-element |

**Supplementary Table 4.** Primers to screen for *P*-element insertions in *CR43651*
